# A community-driven roadmap to advance research on translated open reading frames detected by Ribo-seq

**DOI:** 10.1101/2021.06.10.447896

**Authors:** Jonathan M. Mudge, Jorge Ruiz-Orera, John R. Prensner, Marie A. Brunet, Jose Manuel Gonzalez, Michele Magrane, Thomas Martinez, Jana Felicitas Schulz, Yucheng T. Yang, M. Mar Albà, Pavel V. Baranov, Ariel Bazzini, Elspeth Bruford, Maria Jesus Martin, Anne-Ruxandra Carvunis, Jin Chen, Juan Pablo Couso, Paul Flicek, Adam Frankish, Mark Gerstein, Norbert Hubner, Nicholas T. Ingolia, Gerben Menschaert, Uwe Ohler, Xavier Roucou, Alan Saghatelian, Jonathan Weissman, Sebastiaan van Heesch

## Abstract

Ribosome profiling (Ribo-seq) has catalyzed a paradigm shift in our understanding of the translational ‘vocabulary’ of the human genome, discovering thousands of translated open reading frames (ORFs) within long non-coding RNAs and presumed untranslated regions of protein-coding genes. However, reference gene annotation projects have been circumspect in their incorporation of these ORFs due to uncertainties about their experimental reproducibility and physiological roles. Yet, it is indisputable that certain Ribo-seq ORFs make stable proteins, others mediate gene regulation, and many have medical implications. Ultimately, the absence of standardized ORF annotation has created a circular problem: while Ribo-seq ORFs remain unannotated by reference biological databases, this lack of characterisation will thwart research efforts examining their roles. Here, we outline the initial stages of a community-led effort supported by GENCODE / Ensembl, HGNC and UniProt to produce a consolidated catalog of human Ribo-seq ORFs.

## INTRODUCTION

Human gene annotation is performed by several major reference databases. These resources are used worldwide to support both primary scientific research and clinical workflows, and knowledge gained from downstream applications can be used to improve the annotations in a reciprocal manner. It is now clear that Ribo-seq (also known as ribosome profiling) has the potential to be an important source of biological information. Ribo-seq provides a direct readout of mRNA translation and can therefore nominate translated open reading frames (ORFs) - the precise coding regions of our genomes - with nucleotide resolution. This high-throughput RNA sequencing-based method may herald the biggest paradigm shift in gene annotation since the advent of conventional RNA-seq (**Figure 1A-B**). Consequently, there is a community need for reference annotation of ORF translations identified by this technique. Yet, reference annotation projects lack expertise in this area; they neither generate experimental Ribo-seq data nor develop specific analytical software or workflows. Furthermore, Ribo-seq ORF discovery is a rapidly evolving field that provokes a range of questions on biological interpretation, the answers to which will directly inform the annotation process. As such, collaboration between the annotation databases and experts in the research community will be vital for a successful, accurate, and comprehensive ORF annotation that suits community needs.

**Figure 1:**
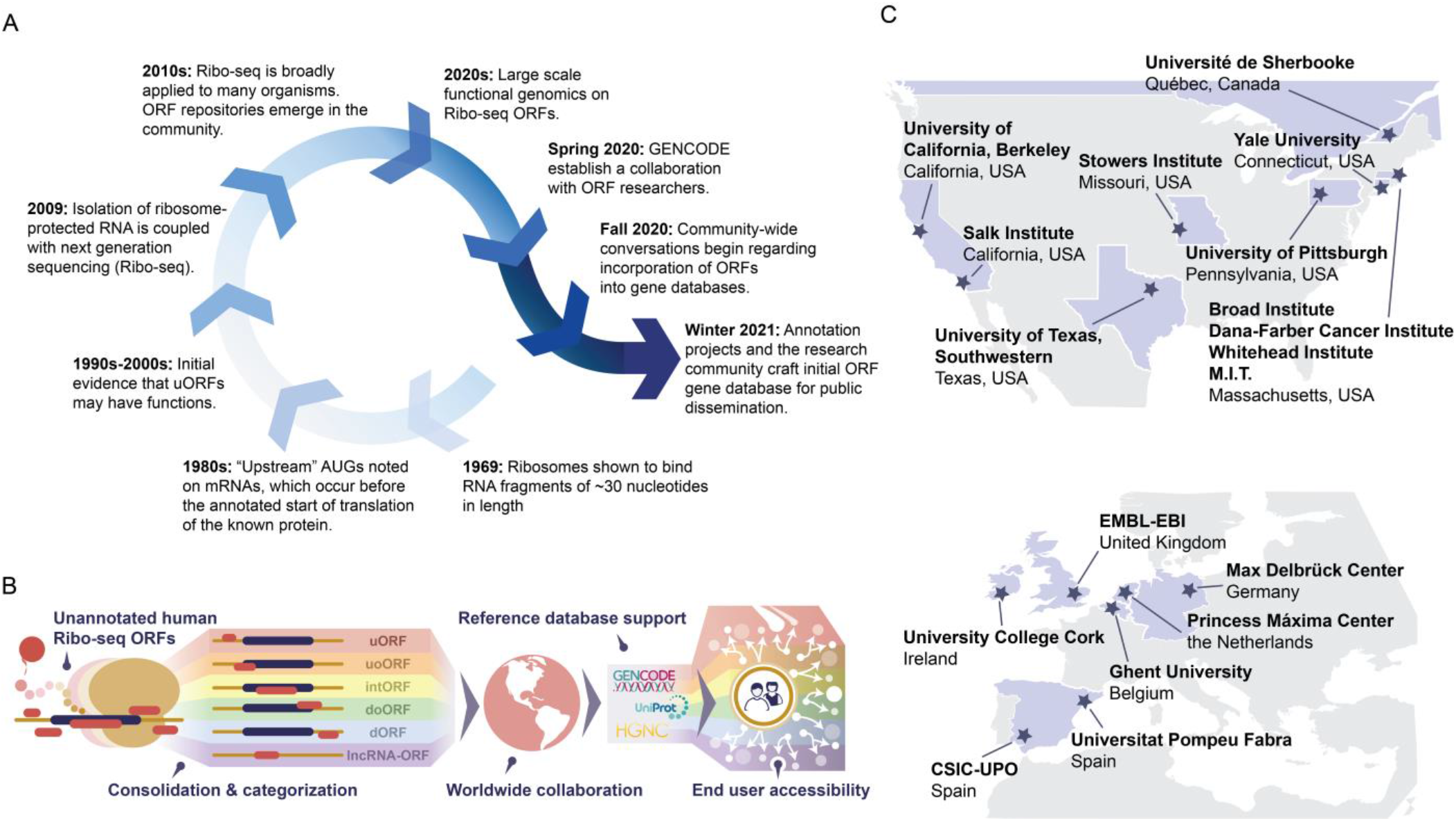
Formation of a community consensus resource for Ribo-seq ORFs. **(A)** A timeline showing the formation of this effort in relationship to major scientific advances in understanding these ORFs. **(B)** A schematic of the main steps done in this effort. **(C)** A map showing the participating institutions included in this effort.

To this end, we have initiated a community-focused collaboration that addresses a central question in modern-day gene annotation: how should one approach Ribo-seq ORF integration? A series of conversations - mediated by Ensembl / GENCODE (henceforth GENCODE) - have been held between frontline Ribo-seq research groups and annotation projects, including UniProtKB / Swiss-Prot (henceforth UniProt) and the HUGO Gene Nomenclature Committee (HGNC). These efforts seek to leverage complementary expertise in the fields of experimental analysis and gene annotation, and reflect a broad, multinational effort including leading experts in diverse domains of gene and protein discovery, genome evolution, Ribo-seq experimental techniques, and computational analysis (**Figure 1C**). In this article, we will discuss the importance of, and also the challenges in, producing a centralized Ribo-seq ORF annotation, and present the first consolidated catalog of Ribo-seq ORFs supported by the major reference gene annotation databases.

### The state of play in coding sequence annotation

A core goal of genome annotation projects is to describe the portion of genomic sequence that encodes protein. It may appear that this goal is close to being accomplished in humans, given that the counts of protein-coding genes in the GENCODE (Frankish et al., 2021; Howe et al., 2021), HGNC (Tweedie et al., 2021), RefSeq (O’Leary et al., 2016) and UniProt (The UniProt Consortium, 2021) gene and protein annotation projects have stabilized at around 20,000. While these gene counts are not static, all four catalogs have incorporated only modest changes in recent years. Furthermore, each project now depends entirely on manual annotation for the addition or removal of protein-coding genes - i.e., the expert human evaluation of evidence available for gene transcription, mRNA translation and protein function - and such adjustments are now made in a largely coordinated manner (Pujar et al., 2018). Nonetheless, new or improved sources of evidence continue to emerge, from both experimental and computational genomics, and these have the potential to instigate new „phases’ in annotation. For example, the recent expansion in availability of genome sequences from other species has greatly improved analytical power in the identification of conserved Coding Sequences (CDS) based on evolutionary methods (Lin et al., 2011), leading to the *in silico* detection of additional protein-coding genes in the last few years (Mudge et al., 2019). Meanwhile, modest numbers of new protein-coding genes continue to be reported via the *de novo* detection of proteins by mass spectrometry (MS) (Ma et al., 2014; Schwaid et al., 2013; Slavoff et al., 2013).

However, such progress conceals an important truth: the annotation of coding sequences remains difficult. While improved evolutionary and MS-based methods continue to be of great value to CDS annotation - especially when used in conjunction - each has its limitations. In particular, evolutionary methods have a potential ‘blindspot’ when it comes to the identification of shorter and/or recently emerged CDSs, e.g. those that are specific to the primate lineage (Ruiz-Orera and Albà, 2019a; Van Oss and Carvunis, 2019). Furthermore, even if the coding evolution of a sequence is supported by conservation and constraint measurements, additional experimental support is required to confirm the production of a protein and elucidate its potential biological role. Similarly, developing CDS annotations exclusively through short peptide fragments obtained by MS is challenging (Nesvizhskii, 2014) and remains a highly extrapolative effort (Wright et al., 2016). The development of methods to obtain „true’ full-length protein sequences would effectively resolve the difficulties of CDS annotation, and while recent advances in proteomics technologies hold promise for the future of protein sequencing (Swaminathan et al., 2018; Timp and Timp, 2020), alternative approaches are required in the meantime.

### Enter Ribo-seq

Given the challenges associated with protein annotation, the experimental datasets produced by Ribo-seq would seem to offer a major boon to annotation projects. First developed in 2009 (Ingolia et al., 2009), Ribo-seq is an experimental method that isolates ribosome-bound RNA fragments for deep sequencing by treating cell lysates with RNase to degrade bulk, free-standing RNA while preserving ribosome-bound RNA fragments of ~30 nucleotides in length. Library preparation and sequencing of these RNA fragments offer genome-wide footprints of ribosome-RNA interactions, enabling the possibility of detecting translated ORFs with sub-codon resolution (Ingolia et al., 2009, 2014; Ruiz-Orera and Albà, 2019a). Thus, in contrast to MS, Ribo-seq maps sequencing reads to the genome or transcriptome, rather than peptide fragments to potential protein sequences.

Collectively, there have been thousands of human genomic sites newly nominated as translated by Ribo-seq, a small subset of which are currently supported by peptide-level evidence in MS data (Bazzini et al., 2014; Chen et al., 2020; Doll et al., 2017; van Heesch et al., 2019; Martinez et al., 2019; Ouspenskaia et al., 2020; Prensner et al., 2021). Many of these regions of prospective translation reside in areas of the genome previously thought to be non-coding, and they can be categorised according to their genomic location and context with regard to existing annotations. **Box 1** outlines the six categories of Ribo-seq ORFs that are considered in this article, all of which are associated with protein-coding genes (uORFs, uoORFs, intORFs, dORFs and doORFs) and long non-coding RNAs (lncRNA-ORFs). Ribo-seq can also describe translations within pseudogenes (Pei et al., 2012; Sisu et al., 2014, 2020), i.e. loci that are currently believed to be defunct protein-coding genes. However, pseudogene translations - ‘pseudo-ORFs’ - are not considered in the efforts outlined here due to potential complexities in mapping Ribo-seq data at loci that can have highly similar paralogs elsewhere in the genome. Although present at much lower numbers compared to other translations, Ribo-seq reads mapped onto circular RNAs (circRNAs) can also be detected, suggesting cap-independent translation (van Heesch et al., 2019; Legnini et al., 2017; Pamudurti et al., 2017; Yang et al., 2017). However, reference annotation projects do not yet incorporate circRNAs, and the experimental evidence that circular transcripts are translated by ribosomes into ‘circORFs’ (or ‘cORFs’) remains a topic of ongoing debate (Hansen, 2020; Ho-Xuan et al., 2020).

#### Box 1: A proposed standardized nomenclature and categorization of Ribo-seq ORFs

We use Ribo-seq ORF to refer to translated sequences identified by the Ribo-seq assay that have not already been annotated by reference annotation projects. Such ORFs have also been referred to as **non-canonical ORFs** or **alternative ORFs (altORFs)** to reflect this lack of annotation, and more recently as novel ORFs (**nORFs**). The terms small ORF (**smORF**) and short ORF (**sORF**) are often used to describe those translations that are under a given size, typically 100 amino acids, while newly-identified small proteins are often referred to as **microproteins**, **micropeptides** or **short ORF-encoded polypeptides (SEPs)**. Ribo-seq ORFs have been traditionally sub-categorised based on their location on the overlapping gene on the same strand, and a variety of nomenclature terms have been used. Our work uses the following six terms.

1. **Upstream ORFs (uORFs)**. uORFs are located within the exons of the 5’ untranslated region (5’ UTR) of annotated protein-coding genes. Many uORFs are known to regulate the translational efficiency of the downstream canonical protein (Johnstone et al., 2016; McGillivray et al., 2018), for example in response to stress (Starck et al., 2016). Others may have roles that are independent from the downstream protein, and some have been shown to produce independent proteins (Cloutier et al., 2020; Huang et al., 2021; Rathore et al., 2018).
2. **Upstream overlapping ORFs (uoORFs).** uoORFs are translated from the 5’ UTR of an annotated protein-coding gene and partially overlap its coding sequence in a different reading frame. At least some uoORFs regulate translation of their overlapping CDS in a manner similar to uORFs but with effects that are anticipated to be stronger, as the ORF terminates within the main CDS (past the CDS initiation codon) (Johnstone et al., 2016). Some uoORFs have been shown to produce independent proteins (Khan et al., 2020; Loughran et al., 2020).
3. **Downstream ORFs (dORFs)**. dORFs are located within the 3’ UTR of annotated protein-coding genes. While ORFs are ubiquitous within 3’ UTRs and can be translated using canonical translation initiation factors (Nobuta et al., 2020), dORFs are less commonly detected in Ribo-seq assays than uORFs, and their putative biological roles remain underexplored. It has been suggested that they can act as *cis* translational regulators (Wu et al., 2020).
4. **Downstream overlapping ORFs (doORFs).** doORFs start within the genomic coordinates of an annotated CDS but their reading frames continue beyond the annotated CDS and terminate in the 3’ UTR of the annotated gene.
5. **Internal out-of-frame ORFs (intORFs).** intORFs - also referred to as **altCDSs**, **nested ORFs** and **dual-coding regions** - are located on the mRNA of an annotated protein-coding gene and are completely encompassed within the canonical CDS. However, they translate via a different reading frame, potentially generating an entirely distinct protein. Detection of intORFs with Ribo-seq is possible but difficult due to the convolution of triplet periodicity signals from two reading frames; it largely depends on the length and translation level of the intORF relative to the overlapping canonical CDS (Erhard et al., 2018; Michel et al., 2012). It is also challenging to estimate the evolutionary constraints acting on intORFs independently from those on the overlapping CDS.
6. **Long non-coding RNA ORFs (lncRNA-ORFs)**. lncRNA-ORFs are encoded by transcripts currently annotated as long non-coding RNAs (lncRNAs), including long intervening/intergenic noncoding RNAs (lincRNAs), long non-coding RNAs that host small RNA species (encompassing microRNAs, snoRNAs, etc), antisense RNAs, and others. Most lncRNAs evolve rapidly (Hezroni et al., 2015) and, accordingly, their translated ORFs often lack strong sequence conservation.

Finally, Ribo-seq can also identify alternative isoforms of annotated proteins, including in-frame N-terminal extensions (e.g. as recently reported for *STARD10* and *ZNF281* (Na et al., 2018)), and internal translation initiation sites that produce shorter proteoforms; the latter especially will be of great benefit to annotation projects as they are very difficult to find through conservation studies.

Over time, a number of public resources have appeared that process and display Ribo-seq datasets for the benefit of the community: sORFs.org, for example, offers a collection of Ribo-seq ORFs under 100 amino acids in size for multiple species based on a standardized data reprocessing platform that includes MS data (Olexiouk et al., 2018), while GWIPS-viz is a genome browser-style interface that displays tracks of publicly available Ribo-seq data aligned to the reference genomes. In contrast, Trips-Viz provides access to Ribo-seq data aligned to reference transcriptomes, and allows detailed analysis including ORF prediction incorporating triplet periodicity and MS data integration (Kiniry et al., 2019; Michel et al., 2018). The OpenProt (Brunet et al., 2019) and nORFs.org (Neville et al., 2020) databases seek to collate experimental support - including Ribo-seq and MS - for *any* ORF that can be extracted from the GENCODE or RefSeq annotation, and so to produce a theoretical catalog of the entire translatome. Furthermore, nORFs.org incorporates information on heritability and selection in human populations. Other projects have focused on specific categories of Ribo-seq ORFs; McGillivray *et al*, for example, have produced a catalog of uORFs with predicted biological activity (McGillivray et al., 2018).

In contrast, Ribo-seq datasets have so far had less impact on the reference human genome annotations produced by GENCODE, RefSeq and UniProt. As noted above, these projects now only annotate new protein-coding genes following manual analysis; they do not incorporate ORF catalogs reported in the literature or by independent databases such as sORFs.org in an unsupervised manner, and they have not developed their own pipelines for processing Ribo-seq data. As a result, few Ribo-seq ORFs have been officially named by the HGNC, (Bruford et al., 2020), who work directly with GENCODE, RefSeq and UniProt. This exclusion of Ribo-seq ORFs from the reference annotation databases has downstream implications: virtually all large-scale gene-based projects, including for example ENCODE (ENCODE Project Consortium, 2012), gnomAD (Karczewski et al., 2020), and the UK BioBank (Van Hout et al., 2020), use reference annotation databases to support their projects. Therefore, such large-scale research efforts pass over Ribo-seq ORFs. Crucially, reference annotation databases also provide the common framework for the interpretation of human variants in the clinic, which means that there is at present effectively no recognised intersection between Ribo-seq ORFs and variation datasets in this setting. In short, while Ribo-seq ORFs remain absent from reference annotation catalogs, they will be harder for everyone - from individual researchers to large consortia - to access and use, limiting comprehensive analysis.

While the focus of the current article is on the annotation of Ribo-seq ORFs identified in human, it should be noted that Ribo-seq has also been used to identify translated ORFs in many other species, such as mouse (Ingolia et al., 2011; Ruiz-Orera et al., 2018), zebrafish (Bazzini et al., 2012, 2014; Chew et al., 2013), fly (Patraquim et al., 2020), nematode (Stadler and Fire, 2011), plant (Hsu et al., 2016; Juntawong et al., 2014), yeast (Blevins et al., 2021; Brar et al., 2012; Carvunis et al., 2012; Durand et al., 2019; Ingolia et al., 2009; Smith et al., 2014; Wilson and Masel, 2011) and bacterial and viral genomes (Finkel et al., 2021; Fremin et al., 2020; Hücker et al., 2017; Stern-Ginossar et al., 2012).

### How to appraise Ribo-seq ORFs

Ultimately, Ribo-seq data represents both an opportunity and a challenge for annotation projects. The advantage of being able to identify missing translated elements – and to validate existing coding annotations – is clear. Nonetheless, there are real difficulties in designing an appropriate annotation framework for these datasets, largely because there are significant questions on the precise biological interpretation of a given translated ORF (**Box 1**). Such ambiguities are in conflict with the remit of reference annotations to provide high confidence interpretations (Mudge and Harrow, 2016). Thus, the hesitancy to annotate Ribo-seq ORFs can also be explained in terms of ‘usability’. Any ambiguities in an ORF annotation catalog would be passed on to the vast numbers of projects that depend on reference gene annotation, and this ‘burden of responsibility’ is a core consideration when strategizing. For example, if Ribo-seq ORFs are incorporated into reference gene annotations, then a framework for the interpretation of putative disease-linked variants found within these features is also required.

A primary concern is that Ribo-seq ORFs would seem to challenge a key principle of gene and protein annotation, that ‘conservation = function’. The majority of Ribo-seq ORFs do not exhibit strong conservation at the amino acid level or signatures of sequence constraints indicative of selection on CDS. Reference annotation projects have historically avoided the annotation of lineage-specific ‘*de novo*’ proteins, unless clear experimental evidence for the existence of the protein has been provided. In addition, Ribo-seq ORFs also challenge the assumption that ‘translation = protein production’. The experimental technique itself cannot distinguish an ORF that produces a genuine protein from an ORF whose translation product is quickly degraded; it only monitors the synthesis event, not what comes afterwards. This allows for a wide breadth of alternative explanations for the biological mechanisms of Ribo-seq ORF function, as outlined in **Box 2**. For example, certain translations - especially within the uORF, uoORF and dORF categories - have been demonstrated to impart *cis* gene regulatory effects arising from the act of translation rather than from the protein that is being synthesized (Andreev et al., 2015; Johnstone et al., 2016; Starck et al., 2016). uORFs in particular have been known to exist for decades - stemming from investigations by Kozak and others about why a CDS does not always employ the first AUG present in a given mRNA (Kozak, 1991; Lovett and Rogers, 1996) - although reference annotation projects have made very little progress in their annotation (**Figure 1A**).

#### Box 2: Interpreting Ribo-seq ORFs

There are multiple possible cellular interpretations of Ribo-seq ORF translations. Below, we list several of the most likely possibilities. Note that a given ORF may encompass several of these possibilities, e.g., a translated ORF that is both regulatory and implicated in disease neoantigen production.

1. **A Ribo-seq ORF encodes a ‘missing’ conserved protein**. Such ORFs can be recognised as canonical – in accordance with existing protein annotations – on the basis that the sequence of the protein they encode shows clear evidence of being maintained by evolutionary selection over a significant period of evolutionary time (Magny et al., 2013; Pauli et al., 2014).
2. **A Ribo-seq ORF encodes a taxonomically restricted protein**. Such ORFs encode proteins whose sequence and molecular activities are specific to one species or lineage. Evidence for selection or conservation across distant species or lineages is lacking for these ORFs, either because the protein sequence has diverged beyond recognition from its orthologues, or because the protein evolved recently from previously noncoding material and homologues do not exist in other species or lineages (Vakirlis et al., 2020a).
3. **A Ribo-seq ORF regulates protein or RNA expression**. Translation of regulatory ORFs does not result in a protein product under selection but regulates the expression of a canonical protein. This paradigm is well established for uORFs and uoORFs, as noted in **Box 1**, though it is potentially applicable to other transcript scenarios. Regulatory ORFs may compete for ribosomes with their downstream canonical ORFs or produce nascent peptides that stall ribosomes (Lovett and Rogers, 1996), leading to the controlled ‘dampening’ of protein expression (Johnstone et al., 2016). Alternative modes of action, such as the induction of RNA decay pathways, the processing of small RNA precursors or the adjustment of RNA stability, have also been inferred (Carlevaro-Fita et al., 2016; Sun et al., 2020; Tani et al., 2013).
4. **A Ribo-seq ORF is the result of random translation**. The translation of some Ribo-seq ORFs may simply be ‘noise’. Since translation has a high bioenergetic cost (Lynch and Marinov, 2015), a protein that results from random translation is likely to be translated at lower levels than a canonical CDS and evolve neutrally (Ruiz-Orera et al., 2018); it may also be unstable in comparison, and be potentially rapidly degraded. Nonetheless, it is theoretically possible that certain proteins do exist as stable ‘junk’ proteins, or that random translation events affect the expression of the canonical protein. The detection of random ORFs is less likely to be reproducible.
5. **A Ribo-seq ORF translates a disease-specific protein.** This protein would not be produced under normal physiological homeostasis but could be of major interest for diagnostics and therapeutics. Insights are especially emerging in cancer biology, where transcription and translation are known to be dysregulated. This leads to the production of ‘aberrant’, possibly rapidly-degraded proteins that are commonly antigenic and presented on the cell surface by the HLA system, offering the prospect of neoantigens (Chong et al., 2020; Laumont et al., 2016, 2018; Ouspenskaia et al., 2020; Ruiz Cuevas et al., 2021). In addition, antigens resulting from disease-specific dysregulated ribosome activity - sometimes called defective ribosomal products (DRiPs) (Rock et al., 2014; Yewdell et al., 1996) - have also been explored.

### New ORFs in the light of evolution

Sequence conservation and constraints on ORF codon structure across species indicate the action of purifying selection maintaining the protein encoded by an ORF. Traditional methods to assess sequence homology and purifying selection are size-dependent and do not work well with Ribo-seq ORFs, which are commonly small (Couso and Patraquim, 2017; Ruiz-Orera and Albà, 2019b). In contrast, recent methods like PhyloCSF (Lin et al., 2011) have single-codon resolution. The recent creation of whole genome PhyloCSF datasets for several species led to the detection of a set of short human proteins conserved within the mammalian order, several of which have been experimentally characterized (Anderson et al., 2015; Cloutier et al., 2020; Mudge et al., 2019; Pauli et al., 2014; Rathore et al., 2018). However, most Ribo-seq ORFs display conservation limited to primates, with no evidence of selective constraints acting to maintain protein sequences (Prensner et al., 2021; Ruiz-Orera and Albà, 2019a).

The absence or weakness of signatures of purifying selection observed in Ribo-seq ORFs could suggest that most do not encode proteins that contribute to organismal fitness (Aspden et al., 2014; Gerstein et al., 2007; Ji et al., 2015; Kaessmann, 2010; Ruiz-Orera and Albà, 2019a). However, purifying selection is unlikely to yield detectable sequence signatures on ORFs that have evolved recently *de novo* from transcribed sequences that were previously non-coding (Van Oss and Carvunis, 2019). Most human-specific and primate-specific proteins have likely evolved through this mechanism (Vakirlis et al., 2020a). ORFs that are so ‘young’ in evolutionary terms may not have existed for a long enough time to display detectable sequence signatures of selection, but they could still be beneficial for fitness (Prensner et al., 2021; Vakirlis et al., 2020b). Our knowledge that ORFs can also mediate gene regulation complicates matters further, especially as less is known about the mode and tempo of regulatory ORF evolution. Such ORFs could potentially be identified by alternative patterns of conservation, assuming that this regulatory effect is not specific to the human lineage. For instance, the nucleotide sequences encoding uORFs may harbor specific conserved secondary structure features that impact their translational efficiency or repressive effect against translation of the CDS (Chew et al., 2016). Alternatively, it may be that the position of a Ribo-seq ORF in a transcript is conserved, i.e. that the initiation and / or termination codons are conserved, or that the length of the reading frame - although not the specific protein sequence - is constrained (Spealman et al., 2018). Numerous uORFs and dORFs have now been identified that are conserved between orthologous genes in vertebrates, despite sharing little similarity at the amino acid level (Bazzini et al., 2014; Chew et al., 2016; Dumesic et al., 2019; Wu et al., 2020). It may be appropriate to consider ORFs with these properties as candidate regulatory elements. However, there can be alternative explanations for the observed conservation of Ribo-seq ORFs at the DNA level. Some intORFs, uoORFs and doORFs will inevitably have elevated conservation due to the presence of selection in the alternative CDS reading frame, while uORFs and dORFs can overlap with conserved non-coding sequence elements including RNA secondary structures, protein-RNA binding motifs and microRNA binding sites (Ruiz-Orera and Albà, 2019b). Altogether, these considerations make it very difficult to infer whether a Ribo-seq ORF contributes to organismal fitness based on its sequence evolution alone.

### A community-driven approach to Ribo-seq ORF annotation

The increasing trove of Ribo-seq data and the growing community interest in newly detected Ribo-seq ORFs have ‘forced’ the issue of their annotation. It is clear that Ribo-seq datasets *do* contain insights of great value for genome annotation. To this end, the GENCODE gene annotation project recently convened a broad panel of experts in the field of RNA translation, ribosome profiling, noncoding RNA biology and ORF detection, in order to begin this work in a ‘community-led’ manner. The major goals of this effort are: (1) to ensure appropriate technical analysis and incorporation of existing Ribo-seq datasets; (2) to establish a consensus strategy for the biological interpretation of Ribo-seq data; (3) to convert this into a systematic framework for gene annotation that is appropriate for wide-ranging public use.

For the research community, this work seeks to solve several existing problems. Primarily, current efforts to find ORFs lack coordination. Although the field is in its infancy, it is clear that ORF collections provided by research publications often have significant overlaps. We anticipate that groups will continue to ‘rediscover’ ORFs that have already been reported while there remains no ‘consensus’ ORF catalog with widespread buy-in from the research community; existing databases such as sORFs.org (Olexiouk et al., 2018), OpenProt (Brunet et al., 2019) and the uORF catalog produced by McGillivray *et al* (McGillivray et al., 2018) employ disparate parameters not agreed upon at the community level, and these catalogues are not embedded in the gene annotation‘ecosystem’.

To this point, it is clear that representation in reference annotation databases maximises visibility, and the importance of this cannot be overstated. Pertinently, there are numerous examples of ORFs that have been investigated independently and in great detail by multiple research groups. For example, *LINC00116* was first reported to produce a 56 amino acids protein in 2013 (Catherman et al., 2013). Subsequently, a series of publications assigned different names to this protein including MTLN (Chugunova et al., 2019; Stein et al., 2018), MOXI (Makarewich et al., 2018), MPM (Lin et al., 2019), and LEMP (Wang et al., 2020) (it is now officially named as protein-coding gene *MTLN*). Another lncRNA example is the murine Gm7325, shown to encode an 84 amino acids protein termed Myomixer (Bi et al., 2017), Myomerger (Quinn et al., 2017) or Minion (Zhang et al., 2017), since named as ‘myomixer, myoblast fusion factor’ (*MYMX*) in humans. While some of this research may have overlapped in time, the usage of disparate names may engender confusion amongst researchers. Protein-coding ORFs within the 5’ UTRs (uORFs) of *SLC35A4* (Andreev et al., 2015), *MIEF1* (Andreev et al., 2015; Brown et al., 2017; Delcourt et al., 2018; Rathore et al., 2018), *ASNSD1* (Cloutier et al., 2020), and *MKKS* (Akimoto et al., 2013) have also been detected and characterized. These four uORFs are now annotated as protein-coding by GENCODE and named by HGNC, and should no longer be reported as ‘novel’ by studies that compare against the latest gene annotations. Moving forward, we anticipate that, if a centralised ORF collection can be agreed upon by the research and annotation communities, then this will help the field move beyond the initial ORF ‘discovery’ phase into placing the emphasis on experimental characterisation.

### Devising a framework for the annotation of Ribo-seq ORFs

The process to incorporate human Ribo-seq ORFs into reference genome annotations is anticipated to occur in two phases. The first phase **(Phase I**) involves the creation of a consolidated list of literature-reported Ribo-seq ORFs (Calviello et al., 2016; Chen et al., 2020; Gaertner et al., 2020; van Heesch et al., 2019; Ji et al., 2015; Martinez et al., 2019; Raj et al., 2016) that have been matched to GENCODE annotations (**Supplementary Tables S1-3)**. Literature sources for human ORFs were selected based on the comprehensiveness of the dataset, specific focus on large-scale ORF detection, and transparency in reporting multiple categories of detected ORFs in the available supplementary files. Thus, although additional human Ribo-seq datasets have been published that do not focus on ORF detection, for the **Phase I** catalog we have not reanalyzed these data. Our efforts here are restricted to collating previously not annotated ORFs from studies that used their own experimental and computational workflows for ORF detection. Furthermore, computational studies that identified ORFs from Ribo-seq datasets already used by others for ORF detection were excluded to avoid redundancy in reporting ORFs from the same source data twice. This **Phase I** catalog of ORFs has been released with this article (see below for discussion and Supplementary Files). We recognise that the relative paucity of Ribo-seq data across many human tissues and cell lines prohibits a biologically comprehensive list of human Ribo-seq ORFs at this time. As this field has only emerged recently, there is currently no standard, widely accepted approach to identify Ribo-seq ORFs. For instance, among the seven datasets selected for inclusion in **Phase I**, there are various ORF discovery pipelines, reference annotations and human genome builds used. Following this, we anticipate a second phase (**Phase II;** see below) that aims for deeper ORF integration into the GENCODE gene annotation and the wider resources of the Ensembl database. In the intervening period, it will be possible to update the **Phase I** catalog as more Ribo-seq datasets are published and gene annotations continue to expand.

To achieve standardization, we defined a set of 8,805 unique Ribo-seq ORF sequences identified within the above-listed seven studies, each of which were fully mapped to Ensembl Release v.101 (August 2020, equivalent to GENCODE v37, see **Methods and Figure 2A**) on genome assembly GRCh38. Because the selected studies applied different minimum ORF length cut-offs and were inconsistent in their reporting of near-cognate ORFs - ORFs employing a leucine or valine initiation codon rather than methionine - we only included cognate Ribo-seq ORFs that were 16 amino acids or longer. Considerations regarding ORF length and near-cognate ORFs are discussed below. Next, we discarded Ribo-seq ORFs that had already been newly annotated as protein-coding or pseudogene in the GENCODE v37 update, as well as any in-frame variations to existing protein-coding isoforms (**Figure 2A, Supplementary Table S4**). In order to mitigate redundancy in Ribo-seq ORF calls, we collapsed instances of multiple isoforms of the same ORF when they shared ≥ 90% of the amino acid sequence, selecting the longest ORF as representative. In total, across all seven datasets, this results in the annotation and inclusion of 7,264 unique cognate ORFs detected in at least one dataset (**Figure 2A-C, Supplementary Tables S2 and S3**).

**Figure 2:**
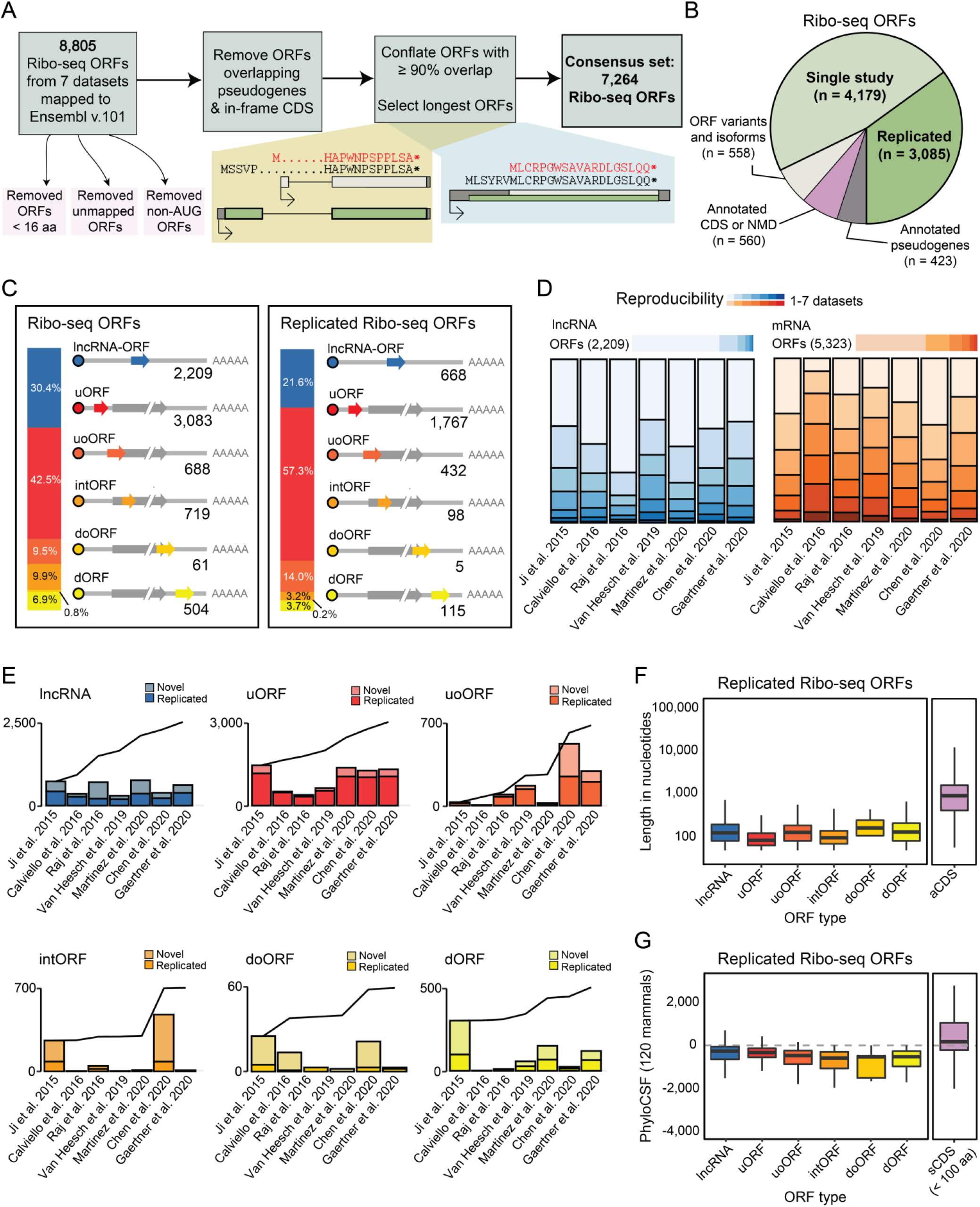
Characterization of a consensus set of Ribo-seq ORFs for annotation by GENCODE. **(A)** A schematic overview of filtration of candidate ribosome profiling (Ribo-seq) ORFs employed in this work. The final consensus list of Ribo-seq ORFs for Phase I includes 7,264 cases. **(B)** A pie chart showing the abundance of ORFs filtered out in each step of the pipeline. **(C)** A diagrammatic representation of all Ribo-seq ORFs (*left*) and a subset of replicated Ribo-seq ORFs (*right*) according to ORF type. **(D)** Replication of Ribo-seq ORFs in lncRNAs (*left*) and in mRNAs (*right*) within each Ribo-seq dataset employed. **(E)** Bar plots with abundances of replicated and non-replicated ORFs across each dataset. Cumulative lines illustrate the evolution in the total number of new unique ORFs identified across studies, sorted chronologically. ORFs are plotted separately for clarity. **(F)** The nucleotide sequence length of replicated Ribo-seq ORFs compared to annotated CDSs (aCDS). ORFs are separated into each respective class. Ribo-seq ORFs are significantly shorter than annotated CDS (two-sided Wilcoxon test, p-value < 10^−10^) **(G)** The phyloCSF scores assessing amino acid purifying selection for ORFs compared to annotated CDSs less than 100 amino acids in length (short CDS, sCDS). Only 2.4% of the replicated Ribo-seq ORFs displayed positive PhyloCSF scores, in contrast to 48% of the sCDS.

### How reproducible are Ribo-seq ORF identifications?

One barrier for GENCODE in assembling a usable Ribo-seq ORF dataset is the variability in ORF identifications. In light of this, one approach to gain confidence in Ribo-seq data is to evaluate how frequently the same ORF is identified across independent studies. This approach has significant technical considerations, due to the fact that the existing studies employ various protocol variations and computational pipelines to process Ribo-seq data. These computational methods (including but not limited to RibORF (Ji et al., 2015), ORF-RATER (Fields et al., 2015) and RiboTaper (Calviello et al., 2016)) can identify ORFs with significant frame preference by additionally evaluating distinct features that reflect the dynamics of translation initiation, elongation and termination (Calviello and Ohler, 2017). For the purposes of **Phase I**, we have used the existing Ribo-seq ORF calls reported in the original manuscript, and we have focused on the list of Ribo-seq ORFs found by more than one dataset as an indicator of robustness of the Ribo-seq signal and low chance that such ORFs reflect spurious variations in data processing methods. These were called “replicated” ORFs (**Figure 2B-E**, **Supplementary Table S2**). In total, we identified 3,085 replicated ORFs: 668 ORFs from annotated lncRNAs (30.1% of all identified lncRNA-ORFs), 1,767 uORFs (57.3% of all uORFs), 432 overlapping uORFs (62.8% of all uoORFs), 115 downstream ORFs (12.8% of all dORFs), 5 downstream overlapping ORFs (8.2% of all doORFs) and 98 internal out-of-frame ORFs (13.6% of all intORFs). Consistent with known features of Ribo-seq ORFs, this set of replicated ORFs was significantly smaller in size compared to annotated CDSs and presented lower PhyloCSF scores - i.e. demonstrated less evidence for evolution as protein-coding sequences - compared to annotated CDSs of a similar small size (**Figure 2F,G**).

It is important to note that reproducibility of these 3,085 replicated ORFs demonstrates consistency in the Ribo-seq signal, but it does not confirm that an ORF has a biological role nor provides insights into what its function may be. Conversely, this selection process does not indicate that the 4,179 ORFs not replicated across multiple studies are “false”. Indeed, Ribo-seq analyses in human cell types remain fairly limited. Since transcription is commonly restricted depending on tissue or cell type, especially for evolutionarily recent genes, we would expect a similar pattern for translation of ORFs encoded by these genes. Moreover, dozens of ORFs are only translated during specific biological processes, such as cell stress (Ramilowski et al., 2020; Starck et al., 2016). Thus, for example, the dataset from van Heesch *et al* is taken only from heart (van Heesch et al., 2019), while none of the other studies include this tissue, and therefore a prejudicial view against any ORFs found in this study alone would be unjustified.

### Consideration of Ribo-seq ORF length

To our knowledge, there is no specific size threshold at which an oligopeptide sequence gains the capacity to be a *bona fide* protein, and reference annotation projects do not have minimal size criteria for protein sequences. Protein secondary structure is thought to be possible at approximately 14 amino acids (Imura et al., 2014) for alpha helices and potentially even less for other secondary structure formations (Kumar and Kishore, 2010), although it is known that some protein sequences are unstructured (Dyson et al., 2005). Even before the advent of Ribo-seq, small proteins had been well characterized in numerous organisms. For example, tarsal-less (*tal*) gene, also named mille-pattes (*mlpt*) or polished-rice (*pri*), produces a polycistronic transcript translated into proteins as short as 11 amino acids in several insect species, such as *Drosophila melanogaster* (Galindo et al., 2007) and *Tribolium castaneum (Savard et al., 2006)*. In humans, *SLN* encodes a muscle-specific 31 amino acid protein known to interact with several Ca(2+)-ATPases reducing the accumulation of Ca(2+) in the sarcoplasmic reticulum (Bal et al., 2012), and this protein has homology extending to fly genomes (Magny et al., 2013). Small proteins may also be medically-important targets: the *CD52* gene encodes a cell surface antigen that is only 61 amino acids but is clinically actionable via the monoclonal antibody alemtuzumab, which has been FDA-approved since 2001 (Dumont, 2001). With the assistance of Ribo-seq and computational approaches such as PhyloCSF, highly conserved human proteins as short as 25 amino acids have already been discovered and annotated prior to our work here (Anderson et al., 2015; Mudge et al., 2019; Pauli et al., 2014). Among the smallest known human protein products are the enkephalin neuropeptides, which are just 5 amino acids in length (Gayen and Mukhopadhyay, 2008; Marcotte et al., 2004), although these are produced by proteolytic cleavage from a larger precursor protein. Among Ribo-seq ORFs, there is limited evidence that even small ORFs can serve as regulatory elements (e.g., *ATF4* uORF1 is 3 amino acids long (Vattem and Wek, 2004), and *AMD1* uORF is 6 amino acids long (Ruan et al., 1996)).

For the purposes of this effort, we have considered ORFs that are at least 16 amino acids in length. Our threshold is largely a practical one, as the literature studies from which we derive our **Phase I** ORFs are inconsistent in their reporting of ORFs smaller than this threshold. Moreover, while peptides derived from proteolytic cleavage may have physiologic roles in some contexts, the process of vetting very small ORFs from Ribo-seq data becomes increasing complicated as computational methods for Ribo-seq data may nominate very short ORFs with variable fidelity, and such ORFs may be too small for unambiguous detection by mass spectrometry due to limitations in aligning MS peptides of very small size. Nevertheless, we recognize that ORFs < 16 amino acids are numerous in Ribo-seq data (**Figure 1B**) and some of them can be replicated in more than one Ribo-Seq dataset (**Supplementary Table S5**), which may prove to be an important area for ongoing investigation and an opportunity for GENCODE to be more inclusive in terms of minimal ORF sizes in the future.

### Near-cognate initiation codons among Ribo-seq ORFs

While the translation of ORFs using an AUG cognate initiation codon is the most frequent and efficient (Kearse and Wilusz, 2017; Kolitz et al., 2009), other near-cognate codons can initiate translation in specific sequence contexts (Diaz de Arce et al., 2018). Generally, CUG is the most efficient near-cognate start codon followed by GUG (Kearse and Wilusz, 2017). One prominent example is the MYC oncogene, which has a well-established CUG start site used in some contexts (Hann et al., 1988), though no obvious phenotypic consequences of this alternative protein product have yet been reported (Blackwood et al., 1994). Computational prediction of near-cognate translational initiation sites (TISs) does not always result in unambiguous identification of TIS positions (Lee et al., 2012). Ribo-seq following lactimidomycin or homoharringtonine treatment can be used to accurately identify TISs; these drugs inhibit translation elongation (Ingolia et al., 2011; Lee et al., 2012) and so result in accumulation of Ribo-seq reads at the putative TIS.

Only four of the seven selected ORF datasets reported near-cognate ORFs, limiting our ability to assess their replicability across studies. As such, we have elected to exclude them from the initial **Phase I** release, though future ORF annotation releases may elect to incorporate them as more data becomes available. For the purposes of this manuscript, we have enumerated the abundance of near-cognate ORFs available in the four datasets reporting them. These studies identified a total 10,412 unique near-cognate ORFs, which we remapped to Ensembl v.101, with CUG (49%) and GUG (19%) being the most frequent non-AUG TISs, as also reported by other Ribo-seq studies (Fritsch et al., 2012; Lee et al., 2012). For several specific cases, we observed near-cognate ORFs that have a strong trinucleotide periodicity signal, do not contain any nearby AUG codon upstream or downstream of the TIS, and are replicated across studies. For instance, *BPNT2* uORF starts with CUG and translates a small ORF of 36 amino acids (**Figure 3A**). However, for ~17% of the near-cognate ORFs, other studies reported a different overlapping translation starting with AUG (e.g., *ANKRA2* uORF; **Figure 3B**), or even different ORFs starting with a mix of alternative near-cognate and AUG codons (e.g. *ZBTB11-AS1* lncRNA-ORF; **Figure 3C**). Such ORFs may participate in cell biology: in the case of *ZBTB11-AS1* ORF, recent experimental data has now contextualized a role in maintenance of cell survival (Prensner et al., 2021). In addition, only ~6% of the near-cognate Ribo-seq ORFs could be replicated in at least 2 of the analyzed Ribo-seq datasets, illustrating the challenge of accurately predicting near-cognate TISs and ORFs in these experiments.

**Figure 3:**
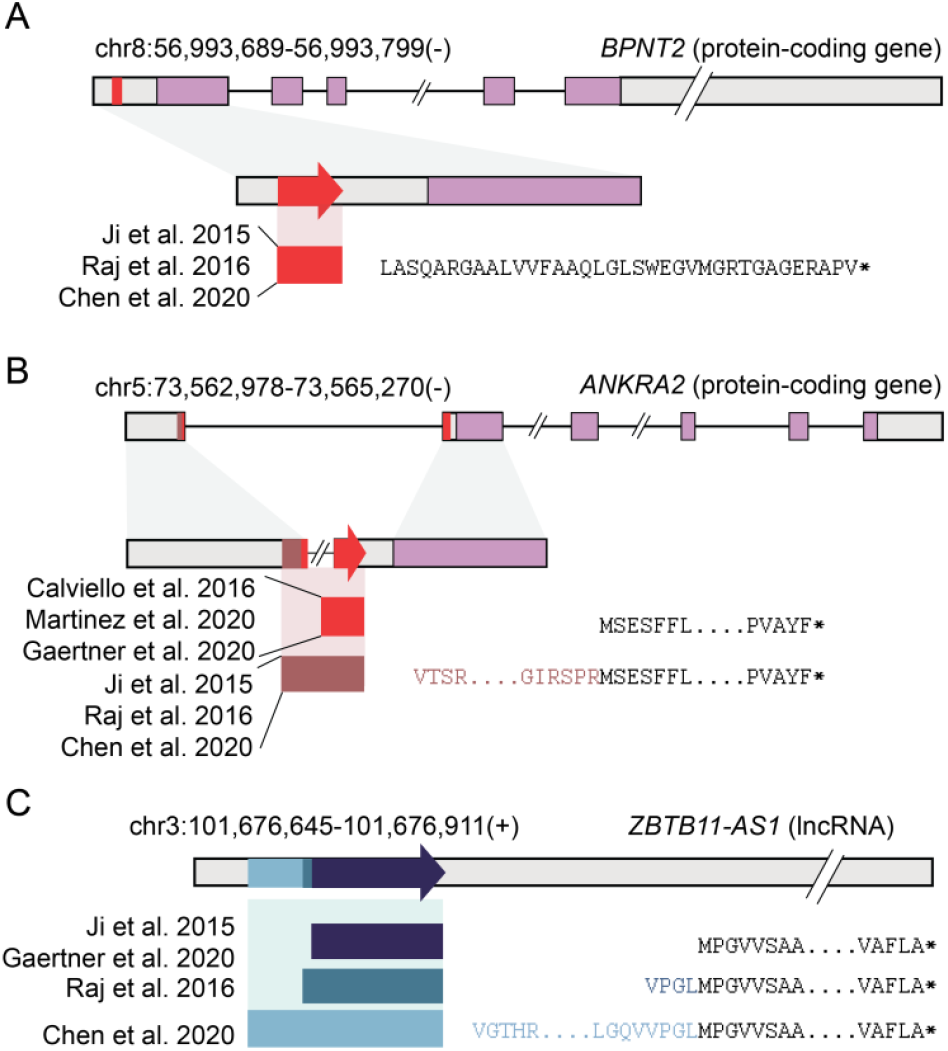
Near-cognate ORFs with consistent or inconsistent Ribo-seq nominations. **(A)** A uORF located in the *BPNT2* gene has multiple datasets supporting translation at a near-cognate initiation codon. **(B)** A uORF located in the *ANKRA2* gene has a near-cognate initiation codon but also a separate annotation utilizing a methionine initiation codon. **(C)** A Ribo-seq ORF located in the *ZBTB11-AS1* lncRNA has several proposed initiation codons utilizing either a methionine initiation codon or a near-cognate initiation codon.

### Conserved protein-coding genes within the Ribo-seq ORF catalog

Ribo-seq ORFs will require detailed, multifaceted approaches to elucidate their biological roles, and a core goal of this work will be to find those that encode bona fide proteins. It is clear that evolutionary metrics cannot be the sole arbitrators of biological function in this regard. Experimental evidence will be especially important for those lineage-specific ORFs that emerged de novo, where their activity at the protein level cannot be inferred by conservation and CDS constraint, and these metrics may be of little value in the study of neo-epitope antigenicity (Laumont et al., 2016, 2018). In a broader context, a myopic focus on protein evolution could also hold back the study of Ribo-seq ORFs that do not impart function at the protein level. This would seem to be especially problematic given the recent insight that purifying selection is widespread across AUG codons initiating uORFs in eukaryotes (Zhang et al., 2021).

Nonetheless, we believe that evolutionary analyses still have a major role to play in Ribo-seq ORF research. Given that the human genome has been vigorously and repeatedly surveyed for missing proteins according to evolutionary profiling in recent years, we might anticipate that very few Ribo-seq ORFs encode ancient proteins. However, as noted, it may be hard to interpret evolutionary signatures in very small CDSs - especially to distinguish CDS constraint from DNA conservation - and so it could be that small proteins commonly ‘fly under the radar’, especially in studies that employ minimal length criteria. Furthermore, we currently lack detailed knowledge as to how algorithms such as PhyloCSF will perform in the identification of dual frame translations, although the protein-coding uoORF recently discovered in *POLG* was able to be identified in this way (Khan et al., 2020). Moreover, a modest number of replicated Ribo-seq ORFs in the datafile *do* show positive PhyloCSF signals (75 out of 3,085 ORFs; 2.4%), potentially indicative of CDS evolutionary constraints.

We have now manually analysed each of these Ribo-seq ORFs according to standard GENCODE annotation criteria, which includes in depth comparative annotation. In total, 10 Ribo-seq ORFs in the datafile have now been annotated as protein coding by GENCODE, as detailed in **Supplementary Table S6**. For clarity, this file lists 15 additional ORFs that have become protein coding in GENCODE within the last two years, following parallel or integrated PhyloCSF and Ribo-seq analyses that predate our current efforts (unpublished data). These annotation decisions have been made unilaterally by GENCODE largely based on the interpretation of PhyloCSF, which demonstrates protein evolution but not protein function, and it will be important to study these translations with additional experimental approaches. We note that the criteria for classifying a protein-coding gene in reference annotation projects does not require certainty in coding status, simply the consideration that this is the most likely explanation of the body of evidence. Furthermore, we do not claim to have identified all conserved proteins in the datafile, and the evolutionary provenance of many Phase I ORFs remains hard to interpret.

We highlight two cases in **Figure 4**. Firstly, a Ribo-seq ORF (*HRURF*) identified within the 5’ UTR of the HR lysine demethylase and nuclear receptor corepressor (*HR*) has been evolving as a protein since at least the base of the mammal radiation (**Figure 4A**). This CDS contains 5 pathogenic ‘non-coding’ ClinVar mutations linked to genetic hair-loss phenotypes that can now be reappraised as missense or loss-of-function protein-coding mutations. These variants were originally noted in a study that identified the ORF but considered it to be a potential regulatory element (Wen et al., 2009); the protein-coding nature of the ORF was not established in annotation catalogs until now. Secondly, we have annotated a Ribo-seq uORF within the highly complex 5’ UTR of ataxin 1 (*ATXN1*) as encoding a protein which is conserved in vertebrate genomes (**Figure 4B**). This is one of five distinct Ribo-seq ORFs from our dataset that are found within the 5’ UTR of this gene; the other four exhibit varying degrees of sequence conservation, although they lack PhyloCSF support for CDS evolution. Interestingly, a nested ORF within the canonical *ATXN1* CDS was previously shown to be translated as a stable protein (Bergeron et al., 2013). Further experimental work will be required to decipher the apparently complex mechanisms of this locus, which is linked to cerebellar ataxias, and recent research indicates that translational control is at least in part mediated by alternative splicing in the 5’ UTR (Manek et al., 2020).

**Figure 4:**
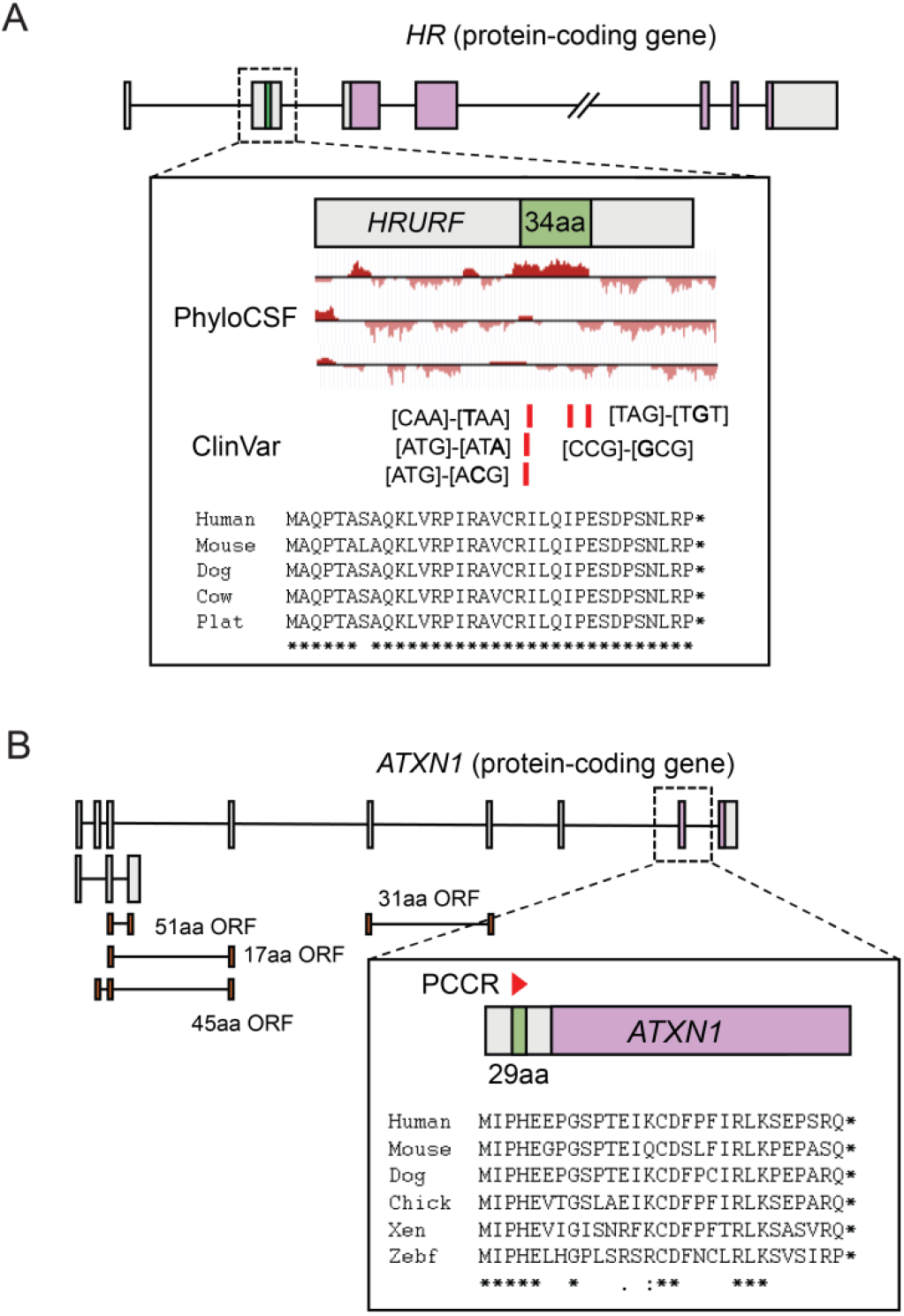
Ribo-seq ORFs in *HR* and *ATXN1* now considered as protein-coding sequences by GENCODE. **(A)** A 34 amino acid uORF (green box) identified within the 5’ UTR of *HR* (UTR sequences in grey boxes; CDS in purple boxes) has been annotated as protein-coding (ENSG00000288677), now recognised as *HRURF* by HGNC. The protein-coding nature of the ORF was inferred by PhyloCSF, according to the positive signal in the top reading frame. Further support was provided by in depth comparative annotation of other vertebrate genomes, demonstrating that the protein likely evolved at the base of the therian mammal radiation; an illustrative alignment is included (‘Plat’ standing for platypus). *HR* has an ortholog in avians and reptiles; the equivalent sequence in these genomes lacks coding potential (not shown), indicating that *HRURF* evolved *de novo*. Five ClinVar variants fall within *HRURF*: RCV000007766.4, RCV001030440.1, RCV000007767.4, RCV000007768.4 and RCV000007769.3 in 5’ order. Each is classed as ‘Pathogenic’, although non-coding. Following the new CDS annotation, mutations RCV000007766.4 and RCV001030440.1 are seen to disrupt the initiation codon, RCV000007767.4 and RCV000007768.4 are missense mutations, while RCV000007769.3 disrupts the termination codon. **(B)** A 29 amino acid uORF within the complex 5’ UTR of *ATXN1* has been annotated as protein-coding, and will appear in a future GENCODE release. This translation has been evolving as coding sequence across the vertebrate radiation (‘Chick’ is chicken, ‘Xen’ is Xenopus, ‘Zebf’ is zebrafish),and the strong PhyloCSF signal (not shown) produced a PhyloCSF Candidate Coding Region (red triangle), indicative of a non-annotated CDS. The canonical transcript of *ATXN1* (ENST00000244769, top model) has six additional 5’ UTR exons with three uORFs inferred from the Ribo-seq datafile (the 17 amino acid and 45 amino acid ORFs are overlapping in different reading frames), while a final ORF has been mapped to an alternatively spliced non-coding transcript (ENST00000467008, second model). While these various UTR exons are generally conserved and supported by transcript evidence in other mammal genomes, the additional ORFs are not strongly conserved and do not present signatures of purifying selection as CDS.

### A resource for Ribo-seq ORFs

Apart from the 10 conserved protein-coding ORFs described in the previous section, GENCODE have not yet integrated Ribo-seq ORFs into the existing ‘data models’ for human protein-coding or lncRNA genes, but present them as a separate datafile in the GENCODE portfolio. These annotations are ‘hygienically’ isolated from the other aspects of the Ensembl resources in order to avoid their unsupervised propagation into downstream projects; users will be able to make the active decision to incorporate them. This does not reflect a general lack of confidence in their potential biological importance, rather our current inability to annotate different modes of ORF function (**Box 2**). In particular, we wish to avoid numerous Ribo-seq ORFs suddenly being included in analyses intended only for validated human protein-coding exons, such as germline or somatic variant calling. The future incorporation of Ribo-seq ORF annotations in the main annotation will require solutions to complex problems in database structure. Many Ribo-seq ORFs are identified on existing mRNA transcripts, i.e. uORFs, uoORFs, dORFs, doORFs and intORFs. Yet it is largely unclear if these scenarios reflect truly polycistronic mRNA translation akin to the polycistronic translation of unique, independently translated proteins observed in viral genomes, and the situation is further complicated by our understanding that translations may instead have a regulatory mode. While it is well established that alternative ORFs on mRNAs can be translated to the latter effect, more information is needed on the potential of the human genome to produce transcripts expressing multiple independent proteins from the same molecule. It could be that distinct proteins are accessed by other mechanisms including alternative transcription. The answers to these questions will have significant practical implications, including in the interpretation of genomic variation. For example, it could be that a given promoter mutation affects the transcription and/or translation of only one of the ORFs. Alternatively, a mutation may affect both ORFs; variants that dampen uORF translation may increase translation of the downstream protein, and conversely variants that increase uORF translation - including gain of function variants that create new uORFs - may dampen downstream translation (Whiffin et al., 2020).

A related question is whether protein-coding Ribo-seq ORFs that are found within the same locus as a known protein - e.g. those in the 5’ UTRs of *HR* and *ATXN1* (**Figure 4A,B**) - should be considered as separate genes. This is not a mere semantic argument. From the biological perspective, there is evidence that some 5’ UTR proteins interact with the downstream known protein (Chen et al., 2020). On the other hand, in several well-studied cases such as the *MKKS* locus (Akimoto et al., 2013) the identified functions of the upstream and downstream proteins appear to be totally distinct. This question is of core importance in reference gene annotation, especially because many users - and in particular large clinical projects - commonly choose to work with one or few transcript models per gene. The ‘visibility’ of protein-coding Ribo-seq ORF annotations is likely to be severely impacted by such approaches if they are merged into the annotation for the known protein, as these short proteins will be at significant risk of being filtered out. For these pragmatic reasons, moving forward, GENCODE, UniProt and HGNC have therefore agreed to annotate protein-coding uORFs, ouORFs, intORFs, doORFs and dORFs as novel genes that are distinct from the known protein at the same locus.

GENCODE decided to be largely agnostic in the **Phase I** ORF catalog, avoiding claims about which ORFs appear more likely to be biologically or medically important, or how they may act. There is little experimental characterization available for most **Phase I** Ribo-seq ORFs, and it is hard to predict which ones will gain additional experimental validation in the future. Therefore, we do not separate ORFs into different ‘biotypes’ (i.e., protein-coding vs. regulatory, etc) as this would be largely speculative at the present time. Instead, we categorize them according to the same schema used in **Box 1** (i.e. uORF, lncRNA-ORF etc). While these specific terms are suitable for our purposes as we develop a unified annotation framework, their usage here is preliminary and does not reflect the development of a finalised classification system.

We recognize the utility of MS in identifying peptide-level evidence for those Ribo-seq ORFs that produce stable proteins. However, MS presents its own set of challenges, and there is significant debate on *whether* or *how* MS methods should be modified to enhance the detection of small ORFs (Chen et al., 2020; Na et al., 2018; Ouspenskaia et al., 2020; Oyama et al., 2004). At the present time there is little consistency between different studies regarding MS data generation and analysis for orthogonal validation of Ribo-seq ORFs. For **Phase I**, we therefore decided to record whether an MS peptide was uniquely identified for a given ORF in its metadata based on the analysis performed in the original study, collecting proposed evidence for 424 replicated Ribo-seq ORFs (Calviello et al., 2016; Cao et al., 2020; Chong et al., 2020; Granados et al., 2014; van Heesch et al., 2019; Kim et al., 2014; Laumont et al., 2016, 2018; Ma et al., 2014, 2016; Martinez et al., 2019; Prensner et al., 2021; Raj et al., 2016; Slavoff et al., 2013; Vanderperre et al., 2013; Zhu et al., 2018) (**Supplementary Tables S2 and S7**). This approach presents the reported data to the user at face value. In the future, **Phase II** will address the topic of mass spectrometry for Ribo-seq ORFs in greater detail.

In summary, our intention is for the ORF catalog produced by **Phase I** of this project to be seen not as the final answer to a problem, rather as a pragmatic interim solution. Moving forward, we will track changes to the Ribo-seq ORF annotations on the GENCODE webportal, and will provide updated versions of the datafile. The annotation of Ribo-seq ORFs considered to be protein-coding will be coordinated between GENCODE, UniProt and HGNC, and also performed in collaboration with RefSeq. Above all, we anticipate that this initial support for Ribo-seq ORFs by reference gene annotation projects will be of benefit to both the research groups discovering and characterising these ORFs, as well as to the broad user community of these biological databases.

### Future perspectives

The recognition of unannotated translation through Ribo-seq data has upended concepts about the translational capacity of the human genome. These data lead to many important questions: How many Ribo-seq ORFs represent canonical protein-coding regions, encoding biologically active protein products? How many Ribo-seq ORFs impart gene regulation through the act of their translation? How do Ribo-seq ORFs arise and to what extent are recently evolved ORFs under selection? What proportion of transcriptional and translational processes that occur in the cell make a minimal contribution to phenotype and fitness? What are the specific implications for Ribo-Seq ORFs in human genetic diversity, health and disease? Will the answers to these questions challenge our definition of a gene, and the current categorical biotyping system employed for gene annotation?

In our view, answering these questions will not be easy, and it is clear that this will not be a short-term endeavour. Furthermore, the development of new biological paradigms will depend on the integration of broad experimental research alongside new analytical methodologies. Already, there have been increasing efforts to dissect the relevance of Ribo-seq ORFs using high-throughput techniques (Chen et al., 2020; Liang et al., 2016; Prensner et al., 2021), genome-wide correlative studies (Calvo et al., 2009; Chew et al., 2016), and individual experimental investigations in diverse contexts (Anderson et al., 2015; Bi et al., 2017; Huang et al., 2021; Pauli et al., 2014). We anticipate that future studies along these lines will allow for further refinement of the ORF catalog, and gene annotations more generally. For example, independently of our work, a 46 amino acid Ribo-seq ORF from our catalog found within the 5’ UTR of *ZNF689* has recently been experimentally validated as a *bona fide* protein, with a prospective *trans* role in transcriptional regulation (Koh et al., 2021).

We also recognize that ongoing Ribo-seq ORF research in other species has the potential to address questions on the evolution of translation, as well as to provide model systems for the experimental investigation of both conserved ORFs and species-specific non-human ORFs (e.g., knock-out mouse models were recently used to investigate a conserved protein-coding uORF in the *PTEN* gene (Huang et al., 2021) and a rodent-specific ORF in the lncRNA *AW112010* (Jackson et al., 2018)). Our roadmap for human Ribo-seq ORF annotation may also provide guiding principles for Ribo-seq ORF incorporation as part of other species annotation projects

Ribo-seq ORFs continue to provoke conversations about gene birth (Keeling et al., 2019; Ruiz-Orera and Albà, 2019a; Ruiz-Orera et al., 2015) and they have opened new vistas for the study of human variation in disease (Whiffin et al., 2020), including variants in these ORFs that may cause human disease phenotypes (Schulz et al., 2018; Wen et al., 2009; Whiffin et al., 2020). It may ultimately be that a significant number of Ribo-seq ORFs prove to be relevant in human health and disease. In light of this, reference annotation databases have the unique capacity to advance this line of research through the propagation and standardization of Ribo-seq ORF datasets, even - and perhaps especially - while the phenotypic impact of these features remains largely uncertain. Establishing a central database of Ribo-seq ORFs will enable greater coordination, consistency and accessibility for researchers globally. Integration of GENCODE efforts with other authoritative resources such as UniProt and RefSeq will further advance these efforts. We envision that the long-term effects of such efforts will yield substantial insight in a key area emerging in human biology.

In this spirit, we hope the results of **Phase I** of this project - as well as our views and considerations on the interpretation and annotation of Ribo-seq ORFs - will be useful and beneficial to the community. We also invite interested labs to reach out and join our future efforts toward the further evaluation and integration of Ribo-seq datasets into new, improved and freely available ORF annotations.

## ACKNOWLEDGMENTS

AF, JMM and PF are supported by the Wellcome Trust [Grant number 108749/Z/15/Z], the National Human Genome Research Institute of the National Institutes of Health under award number 2U41HG007234 and the European Molecular Biology Laboratory. For the purpose of open access, the author has applied a CC BY public copyright licence to any Author Accepted Manuscript version arising from this submission. The content is solely the responsibility of the authors and does not necessarily represent the official views of the National Institutes of Health. Ensembl is a registered trademark of EMBL. MG and YTY are supported by National Human Genome Research Institute of the National Institutes of Health under award number 2U41HG007234. UniProt is supported by the National Human Genome Research Institute (NHGRI) of the National Institutes of Health under Award Number [U24HG007822], European Molecular Biology Laboratory core funds and the Swiss Federal Government through the State Secretariat for Education, Research and Innovation SERI. RP is supported by the Harvard K-12 in Central Nervous System tumors (5K12 CA 90354-18) and the Dana-Farber Trustee Science Committee Award. TFM is supported by the National Institutes of Health under award number F32GM123685. MMA acknowledges funding from the Spanish Government grant PGC2018-094091-B-I00 (MCI/AEI/FEDER,EU) and AGAUR grant 2017SGR01020. JC is supported by the National Institutes of Health Pathway to Independence Award (R00 GM134154) and the Cancer Prevent and Research Institute of Texas (RR200095). JSW is supported by HHMI. AAB is supported by the Stowers Institute for Medical Research and the US National Institutes of Health (R01 GM136849). ARC is supported by funds provided by the Searle Scholars program and the National Institute of General Medical Sciences of the National Institutes of Health award number DP2GM137422. PVB wishes to acknowledge the support from Investigator in Science Award [Grant number 210692/Z/18/Z] by SFI-HRB-Wellcome Trust Biomedical Research Partnership and from Russian Science Foundation [Grant number 20-14-00121]. NH is recipient of an ERC advanced grant under the European Union’s Horizon 2020 research and innovation programme (grant agreement n° AdG788970). NH is supported by a grant from the Leducq Foundation (11 CVD-01). The work of the HGNC is funded by the Wellcome Trust (208349/Z/17/Z) and the National Human Genome Research Institute of the National Institutes of Health (under Award Number U24HG003345). XR and MAB are funded by the Canadian Institutes of Health Research PJT-175322. XR is funded as Canada Research Chair in functional proteomics and discovery of novel proteins. NTI is supported by the U.S. National Institutes of Health under award number R01 GM130996.

## ADDITIONAL DATA

All analyses in this study are performed using published and publicly available analytical tools or software packages. Published Ribo-seq ORF datasets and processed mass-spectrometry peptide datasets were retrieved from the Supplementary Material of each referenced study as described in **Supplementary Tables S1 and S7**. The code used for generating the list of Phase I ORFs is available at https://github.com/jorruior/gencode-riboseqORFs.

**Supplementary File 1:**

**Table S1.** Description of the 7 human Ribo-seq datasets used for this study

**Table S2.** Table with 3,085 replicated Ribo-seq ORFs that were found in at least two Ribo-seq studies. LncRNA-ORFs were divided into two categories (see ORF biotype): lncRNA (lncRNA-ORFs in lncRNA genes) and processed_transcript (lncRNA-ORFs in protein-coding genes)

**Table S3.** Table with 4,179 Ribo-seq ORFs that are from a specific Ribo-seq study. LncRNA-ORFs were divided into two categories (see ORF biotype): lncRNA (lncRNA-ORFs in lncRNA genes) and processed_transcript (lncRNA-ORFs in protein-coding genes)

**Table S4.** Table with 957 Ribo-seq ORFs currently annotated as protein-coding (CDS or NMD) or overlapping pseudogenes in Ensembl v.101. ORFs partially or totally overlapping in-frame CDS were included in this table. If two or more ORFs shared >= 90% of the amino acid sequence, only the longest one was included.

**Table S5.** Table with 1,520 Ribo-seq ORF sequences discarded due to the short size (< 16 amino acids). These ORFs were replicated in at least two Ribo-seq studies.

**Table S6.** This sheet lists 10 ORFs that have been annotated as protein-coding by GENCODE as part of this work, and a further 15 that were previously annotated as part of preliminary investigative work into Ribo-seq datasets combined with in-house PhyloCSF analysis. Proteins that are listed as appearing in GENCODE v38-39 are not in a public ‘genebuild’ release at the time of publication, and so gene and transcript IDs are not yet available for these models. Note that while the ‘comments’ provide brief explanations for the annotation, they do not attempt to establish provenance for the initial identification of the ORF or protein. Further support for these annotations could potentially be found in additional resources or publications. These annotation decisions were made by GENCODE according to an interpretation of the balance of probability when considering all available evidence, i.e. these ORFs are considered *most likely* to be protein-coding, in line with standard annotation criteria. GENCODE recognise that further experimental characterization will be required to support these annotations.

**Table S7.** Description of the 16 mass-spectrometry datasets used for this study. Identified peptides were retrieved from supplementary data and re-mapped to Ribo-seq ORFs.

**Table S8.** Table with 80 Ribo-seq ORF sequences that could not be mapped to any of the current transcript annotations in Ensembl v.101. These ORFs were replicated in at least two Ribo-seq studies.

## METHODS

### Phase I ORF retrieval and mapping

We selected seven ORF datasets from different human studies that represent key projects for genome-wide Ribo-seq ORF identification in the last five years (**Supplementary Table S1**). We retrieved ORF exonic coordinates -when available- and ORF sequences, collecting a total set of 39,788 translated ORFs corresponding to 29,373 unique protein sequences. Only Ribo-seq ORFs found in long non-coding RNAs (lncRNAs), alternative protein-coding frames and/or UTRs from protein-coding genes were selected, discarding annotated proteins, extensions/truncations of annotated isoforms, pseudogenic ORFs and circRNA ORFs. Each of the selected studies applied different minimum length cut-offs to define their Ribo-seq ORFs and only 4 of the studies considered near-cognate ORFs (ORFs starting with non-AUG initiation codons, see **Box 3** and **Supplementary Table S1**). Hence, in order to maximize ORF replicability across studies we discarded 8,503 Ribo-seq ORFs shorter than 16 amino acids and 10,412 Ribo-seq ORFs starting with near-cognate codons.

Next, for the five ORF datasets that were built on an older Human Genome assembly (GRCh37/hg19, **Supplementary Table S1**), we converted ORF coordinates to GRCh38/hg38 using UCSC Liftover (Navarro Gonzalez et al., 2021). We remapped all translated ORF sequences to Ensembl Release v.101 transcriptome (August 2020, equivalent to GENCODE v37), generating a set of 8,805 unique ORFs after excluding 1,767 ORFs that could not be fully mapped to any transcript in this release. We note that the latter set includes 80 replicated Ribo-seq ORFs, i.e., ORFs detected in more than one study (**Supplementary Table S8**); GENCODE are currently examining these Ribo-seq ORFs and the potential transcript structures they would map to for potential inclusion. For 130 ORFs, the sequence could be mapped to more than one transcribed genomic region and the exact unique region was identified by combining the exonic coordinate data and/or the associated gene names annotated in each study. Moreover, we further excluded ORFs overlapping pseudogenes (423) or in-frame coding (CDS) sequences from protein-coding or nonsense-mediated decay transcripts (560) in the current transcriptome version, since the ORF datasets used older transcriptome releases and new protein-coding sequences and pseudogenes are newly annotated in Ensembl v.101 (**Supplementary Table S4**). Finally, in order to get a non-redundant list of translated ORFs for the Phase I, we adapted the clustering method of UniRef90 (UniProt, (The UniProt Consortium, 2021)) and collapsed overlapping ORFs with alternative start or stop codons in groups where multiple instances of ORF variants shared >= 90% of the linear amino acid sequence, considering the longest ORF as representative. If an ORF variant exhibited significant similarity to two or more non-collapsed ORFs, the variant was multiply assigned to all possible cases. This resulted in a total of 7,264 collapsed Ribo-seq ORFs (**Figure 2A**), where only 549 of the ORFs had more than one ORF variant –a total of 558 unique ORF sequence variants. The number of considered ORFs per study varied between 846 and 3,062. This substantial difference in detected ORFs is not necessarily a reflection of data quality or depth, but primarily the result of approach-specific filtering presets or the number of replicate identifications required within each study.

For the **Phase I**, we selected a subset of 3,085 replicated Ribo-seq ORFs that are robustly translated in more than one study. A Ribo-seq ORF was considered as translated in a specific dataset if the main ORF sequence or any of the collapsed ORF variants was found in the ORF list generated by that study.

### ORF classification and transcript assignment

Ribo-seq ORFs were classified into 6 different biotypes defined by the host transcript biotype and the relative position compared to annotated canonical protein-coding sequences (see **Box 1**). However, gene annotations usually contain several overlapping isoforms and 65.44% of the Ribo-seq ORFs could not be unambiguously mapped to a unique host isoform. For these cases, we assigned the transcript with the highest APPRIS score (Rodriguez et al., 2018) as the most likely isoform that translates each ORF. If more than one transcript shared a similar APPRIS score, we further evaluated the Ensembl transcript support evidence (TSL) score. For 1,513 ORFs, the sequence could be still mapped to several transcript sequences with equal support evidence. However, for these cases the selection of different isoforms did not affect the assigned ORF biotype or the exonic coordinates and we randomly selected one of the transcripts as host. All possible Ensembl transcript and gene IDs compatible with each ORF sequence, as well as the IDs of the selected host transcripts, are described in **Supplementary Tables S2 and S3**.

### Multiple-species alignment and PhyloCSF

We downloaded previously generated multiple-genome alignments for 120 mammals (Hecker and Hiller, 2020) and we extracted aligned regions for each of the human Ribo-seq ORFs. Only species where the ORF region could be fully aligned were included in each alignment. As a result, 99.5% and 97.23% of the ORFs were aligned to at least one primate species and a non-primate mammalian species respectively. Next, we assessed the conservation of replicated Ribo-seq ORFs by comparing the patterns of codon evolution in different mammalian species using PhyloCSF (Lin et al., 2011) with default parameters. For comparison, we additionally built multiple alignments and ran PhyloCSF for a set of 531 annotated CDS sequences shorter than 100 amino acids, taking the longest CDS per gene (sCDSs).

### Analysis of proteomics datasets

We searched for additional evidence of Ribo-seq ORF protein production by collecting 16 published datasets that identified peptides mapping to non-annotated protein-coding regions using different mass-spectrometry (MS) approaches (e.g. standard MS/MS, peptidome isolation, HLA peptidome isolation and targeted SRM proteomics, **Supplementary Table S7**). Peptides were retrieved from the corresponding Supplementary Materials and were remapped to the full set of Ribo-seq ORF sequences. Peptides that uniquely mapped to a Ribo-seq ORF and did not map to any annotated protein-coding sequence were retained as MS evidence.

